# On-Demand Fully-Enclosed Superhydrophobic-Optofluidic Device Enabled by High Precision Microstereolithography

**DOI:** 10.1101/2022.06.21.497079

**Authors:** Yu Chang, Mengdi Bao, Jacob Waitkus, Haogang Cai, Ke Du

**Affiliations:** Department of Mechanical Engineering, Rochester Institute of Technology, Rochester, New York 14623, United States; Tech4Health Institute and Department of Radiology, NYU Langone Health, New York, New York 10016, United States; Department of Microsystems Engineering, Rochester Institute of Technology, Rochester, New York 14623, United States; School of Chemistry and Materials Science, Rochester Institute of Technology, Rochester, New York 14623, United States; College of Health Science and Technology, Rochester Institute of Technology, Rochester, New York 14623, United States

**Keywords:** Optofluidics, Microstereolithography, Superhydrophobicity, CRISPR sensing

## Abstract

Superhydrophobic surface-based optofluidics have been introduced to biosensors and unconventional optics with unique advantages such as low light loss and power consumption. However, most of these platforms were made with planar-like micro- and nano-structures, which may cause bonding issues and resulting in significant waveguide loss. Here, we introduce a fully-enclosed superhydrophobic-based optofluidics system, enabled by a one-step high precision microstereolithography procedure. Various micro-structured cladding designs with a feature size down to 100 μm were studied and a “T-type” overhang design exhibits the lowest optical loss, regardless of the excitation wavelength. Surprisingly, the optical loss of superhydrophobic-based optofluidics is not solely decided by the solid area fraction at the solid/water/air interface, but also the cross-section shape and the effective cladding layer composition. We show that this fully-enclosed optofluidic system can be used for CRISPR-labeled quantum dot quantification, intended for in vitro and in vivo CRISPR therapeutics.

## INTRODUCTION

Liquid-core optofluidic configurations, as one of the optofluidic systems, typically consist of an aqueous-core layer and a solid cladding layer^1^. The unique structure of these platforms enable a single miniaturized device to realize liquid sample holding and optical guiding functions simultaneously for applications such as fluorescence-based biosensors^2,3^, absorption spectroscopy^4,5^, and medical studies^6,7^. To achieve an efficient light guiding function, the refractive index of the core layer should be higher than the cladding layer to fulfill total internal reflection (TIR^8^). The principle of TIR is when the incident light goes from the core layer to the cladding layer with an incident angle that is less than the critical incident angle, the light would be totally reflected. There are two ways to reduce the optical loss: one is to choose the appropriate materials of core and cladding layers to maximize the difference of the refractive indexes^9^; The other option is to utilize a complex cladding layer^10^. The latter method is more robust as the refractive index of the analyte is typically low and cannot be easily tuned.

Recently, superhydrophobic surface-based waveguide platforms have been introduced to trap air in the micro-and nanostructures, thus reducing the effective refractive index at the interface^11^. Jonáš et al. established a tapered optical fiber waveguide coupled with a droplet standing on a superhydrophobic surface to measure the quality factors of individual optical resonances^12^. The superhydrophobic surface was made by spin-coated silica nanoparticles on a glass slide. However, the uniformity of such coating is difficult to control. Alternatively, high hydrophobicity was achieved by selective laser ablation on magnesium-fluoride substrates^13^. However, both systems are planar-like, which limit the flexibility to vary the morphology. Previously, we developed a prototype by using sharp-tip black silicon nanostructures as a cladding layer^14^. A low loss of 0.1 dB/cm was achieved with zero incident angle, indicating that the air pockets created by the superhydrophobic nanostructures can reduce the waveguide loss. However, the black silicon nanostructures were created by conventional deep reactive ion etching on a flat and rigid substrate, causing problems for the chip packaging. Similarly, changing the morphology of the nanostructures is extremely challenging with plasma etching, preventing us to study the TIR at the interface.

In this work, high precision microstereolithography^15^, a promising technology to construct complicated 3-D structures with controllable morphology, is used to build a fully-enclosed superhydrophobic-based liquid-core optofluidic device. The microstructures coated with Polytetrafluoroethylene (PTFE) serve as the hydrophobic layer responsible for forming the complex cladding layer and containing air within the microstructures. The solid fraction ratio of the designed cladding layer, cross-section geometry, effective cladding composition, and excitation wavelength are studied to understand the mechanism. This work results in an optimized design guideline for the superhydrophobic cladding optofluidic platforms and is used here for quantum dot-labeled CRISPR Cas-12a biosensing.

## MATERIALS AND METHODS

### Micro-structured optofluidic design

To create a fully-enclosed optofluidic chip with microstereolithography, ultraviolet (UV) curable resin was chosen based on its high resolution and transparency. We created four different micro-structures to study optical loss with a feature size down to 100 μm. Practically, the more air gaps between the created micro-structures, the ratio of liquid-solid contact can be lowered to form a hydrophobic layer. Periodic micro-gratings and micro-pins are shown in **Figure 1a** and **1b** respectively, with aspect ratios of ~5.7 and 8 respectively. These periodic structures have been widely used for various superhydrophobicity applications^11,16^. **Figure 1c** shows a “T-shape” double over-hang design with super-repellent capability since the low contact angle prevents the liquid from entering the air gap^17^. A slightly modified “umbrella” design is shown in **Figure 1d**, which changes the flat “T-shape” cap to a 13.31° slope. The hydrophobicity was fulfilled by the inclined roof structures and the air gaps between them. Next, the microstructures were dip-coated with PTFE (Teflon AF 6%, Chemours) followed by baking at 110□ for 3 hr to create the superhydrophobic surfaces^19^. The thickness of the coating is ~3 μm which was measured by a profilometer (Tencor P2). The cross-section of a liquid filled “T-shape” structure is shown in **Figure 1e**. Air is sustained between the microstructures without wetting. Before coating with PTFE, the material is hydrophilic with a static contact angle of 58° (**Figure 1f**). After coating, the surface exhibited superhydrophobicity with a contact angle of 120° (**Figure 1g**). The ultra-high hydrophobicity that combines the superhydrophobic surface and the overhang structures creates an air gap between the microstructures with a center-to-center distance of 2.8 mm (**Figure 1h**), which is far greater than the distance in optofluidic design (70 μm). The droplet could be “repelled” because of the considerable vertical component of the surface tension on the overhangs. The low refractive index of the air layer creates a light guiding function. As shown in **Figure 1i**, TIR at the solid/water/air interface is achieved, as denoted by the yellow arrow, where light is reflected back into the liquid core with an incident angle of ~35°. As a fully-enclosed optofluidic system, the liquid inlet and outlet were created during the printing step, thus avoiding leaking issues and forming a working device as shown in **Figure 1j**.

**Figure 1.**
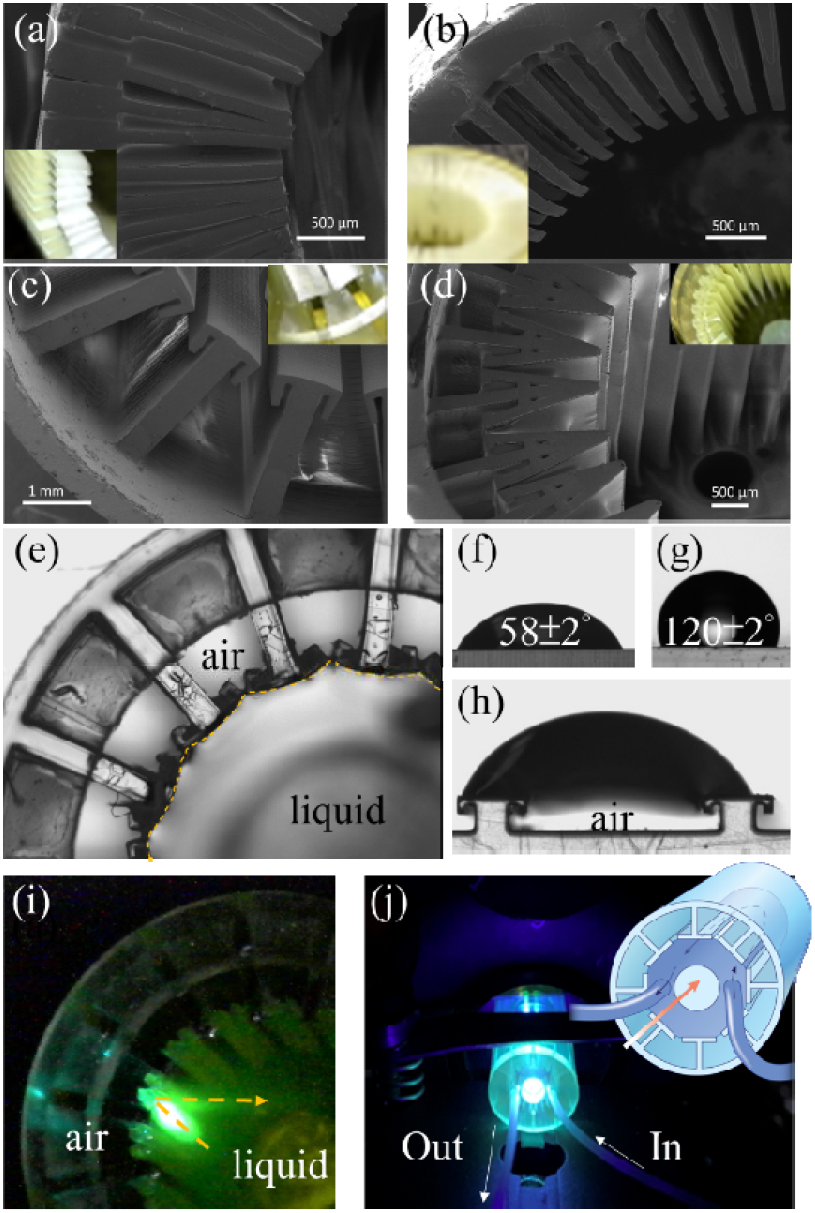
SEM image and optical micrograph of (a) Micro-grating structures. (b) Micro-pins structures. (c) “T-shape”. (d) “Umbrella”. (e) Cross-section of “T-shape” optofluidic device filled with water in the core. The solid/water/air interface is labeled with a dashed yellow line. The static water contact angle measurement of a flat sample (f) before and (g) after Teflon AF coating. (h) The droplet supported by two T-shape structures with a wide gap. (i) Light beam reflected at the solid/water/air interface with an incident angle of 35°. (j) Micrograph of a working optofluidic platform with the light sharing the same path with the liquid core. Liquid is pumped into the liquid core through the embedded micro-tubing. The fluid flow directions in the device are depicted in the inset.

### Transmission measurements

The measurement setup is shown in **Figure 2a**. An optical fiber-coupled LED laser (emitter diameter: 1mm, Thorlab M00559094) is aligned with the optofluidic chip, which is placed on a 3-D stage (Thorlabs MBT616D). The fiber is connected to a UV lens where the incident angle can be tuned. The outlet of the optofluidic chip is connected to a filter (FB610-10) and another optical fiber to collect the light signal and send it to a spectrometer (FLMS16493). The incident angle was fixed at 5° (max transmission intensity) for all samples and tested for four different excitation wavelengths: 405 nm, 490 nm, 595 nm, and 1,100 nm.

**Figure 2.**
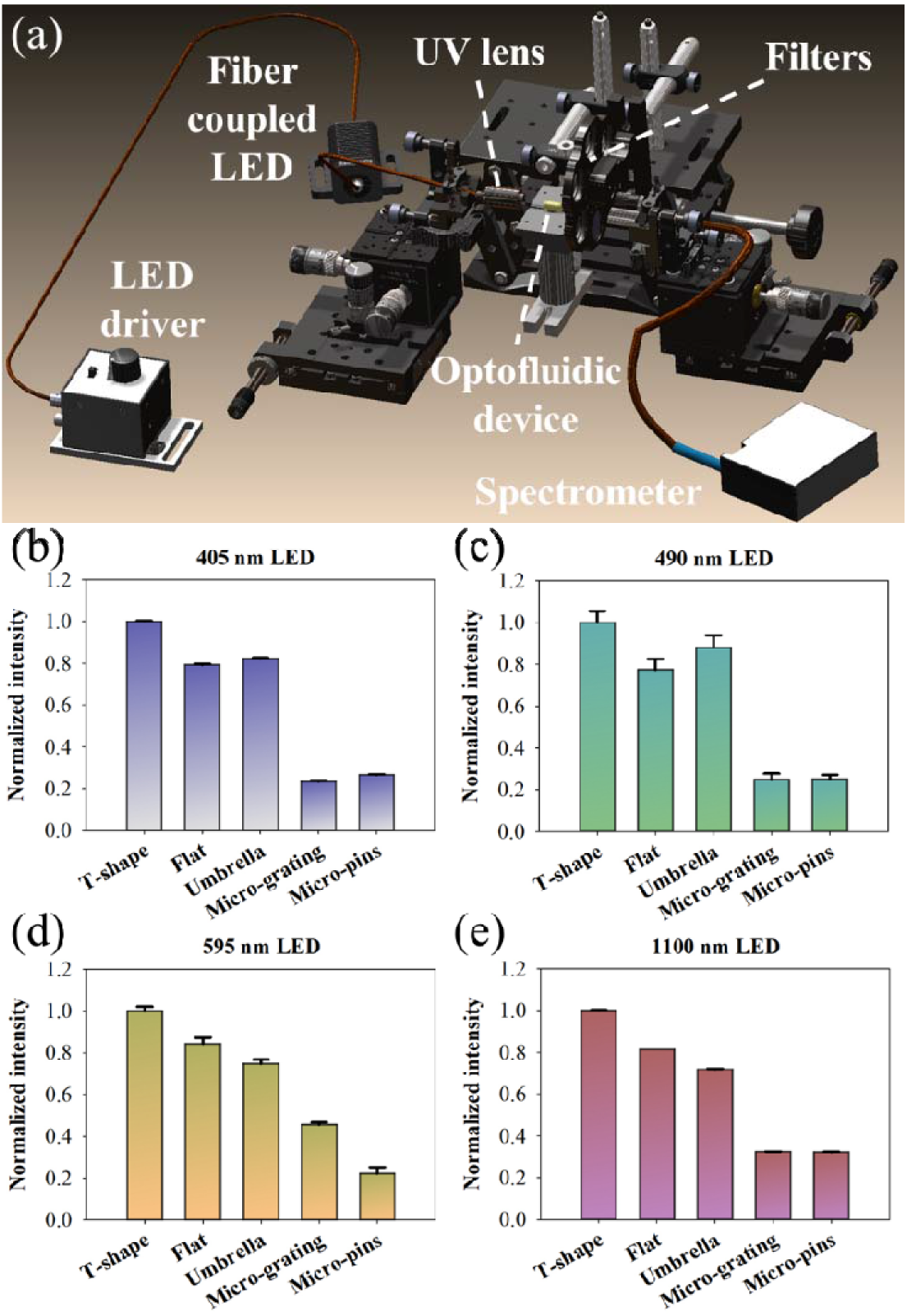
(a) The schematic of the setup for optical measurements. Normalized light transmission of different optofluidic chips with an excitation at (a) 405 nm. (b) 490 nm. (c) 595 nm. (d) 1,100 nm. The error bars represent the standard deviation for each respective sample.

### CRISPR-Cas12a Sensing with the Optofluidic Device

The CRISPR-Cas complexes were first prepared by mixing 1 μL of binding buffer, 62.5 nM of biotinylated single-stranded DNA probes, and 9.5 μL of nuclease free water. After adding the target DNA, the mixture was immediately activated and incubated at 37°C for 2 hr. Next, the reaction product was mixed with 1.68 nM of streptavidin-coated quantum dots at room temperature for 30 min. Following that, the mixture was incubated with anti-fluorescein-coated magnetic beads at room temperature for 30 min, allowing the intact probe-quantum dot conjugate to react with magnetic beads. The newly formed magnetic bead-probe-quantum dot conjugates were then isolated and removed from the desired supernatant which contained the quantum dots with degraded probes due to the CRISPR-Cas complex. Finally, the supernatant was collected, diluted, and added into the optofluidic device.

## RESULTS AND DISCUSSION

As shown in **Figure 2b-e**, for all the measurements, the “T-shape” sample shows the highest transmission, regardless of the excitation wavelength. The flat sample (without any microstructures) shows ~20% lower transmission of the light beam than the “T-shape” sample, followed by the “umbrella” sample, the micro-gratings, and the micro-pins. Based on the 3-D ray tracing analysis^20^, the cladding layer thickness should be smaller than 15% of the core diameter to ensure the light rays travel primarily within the core. As the diameter of the core is 6 mm, we calculated that the thickness of the cladding layer required is 900 μm, theoretically. The calculated solid fraction are, 29.6% for micro-gratings, 42% for micro-pins, 41.2% for flat surface, 86.4% for “umbrella”, and 36.9% for “T-shape”. Since Teflon is very thin compared to the printing material, it is neglected during calculation. Note that for the flat optofluidic device, the solid substrate is solely the 400 μm wall within the required cladding thickness of 900 μm. Based on the theory of TIR, a lower solid fraction for the cladding layer can reduce the optical loss. Thus, we expected the micro-gratings should provide the highest transmission, followed by the “T-shape” and flat samples. As shown in **Figure 2**, “T-shape” and flat samples always show the highest transmission among all the designs, which agree with our theoretical prediction. However, the measured transmission for micro-gratings and micro-pins are significantly lower. This is a result of the microstructures collapsing following the PTFE coating^21^. On the other hand, the mechanical stability of “T-shape” and “umbrella” microstructures are much higher, thus providing improved containment of the liquid in the core while showing better optofluidic performance.

We further study the light guiding capabilities of the microstructures based on the solid cladding thickness, light refraction, and macro scattering. **Figure 3a** shows the transmission intensity of a flat optofluidic chip with a wall thickness of 400 μm and 600 μm, respectively. A 200 μm increase of the wall thickness significantly reduces the light transmission. With a zero incident angle, the light transmission for the 600 μm sample is ~4 times less than that of the 400 μm sample because the solid fraction in the effective cladding layer absorbs the light. To reduce the optical loss, the solution, as described above, is to reduce to wall thickness which is why the 400 μm thick flat optofluidic chip demonstrates good performance even without micro-structure aided superhydrophobicity.

**Figure 3.**
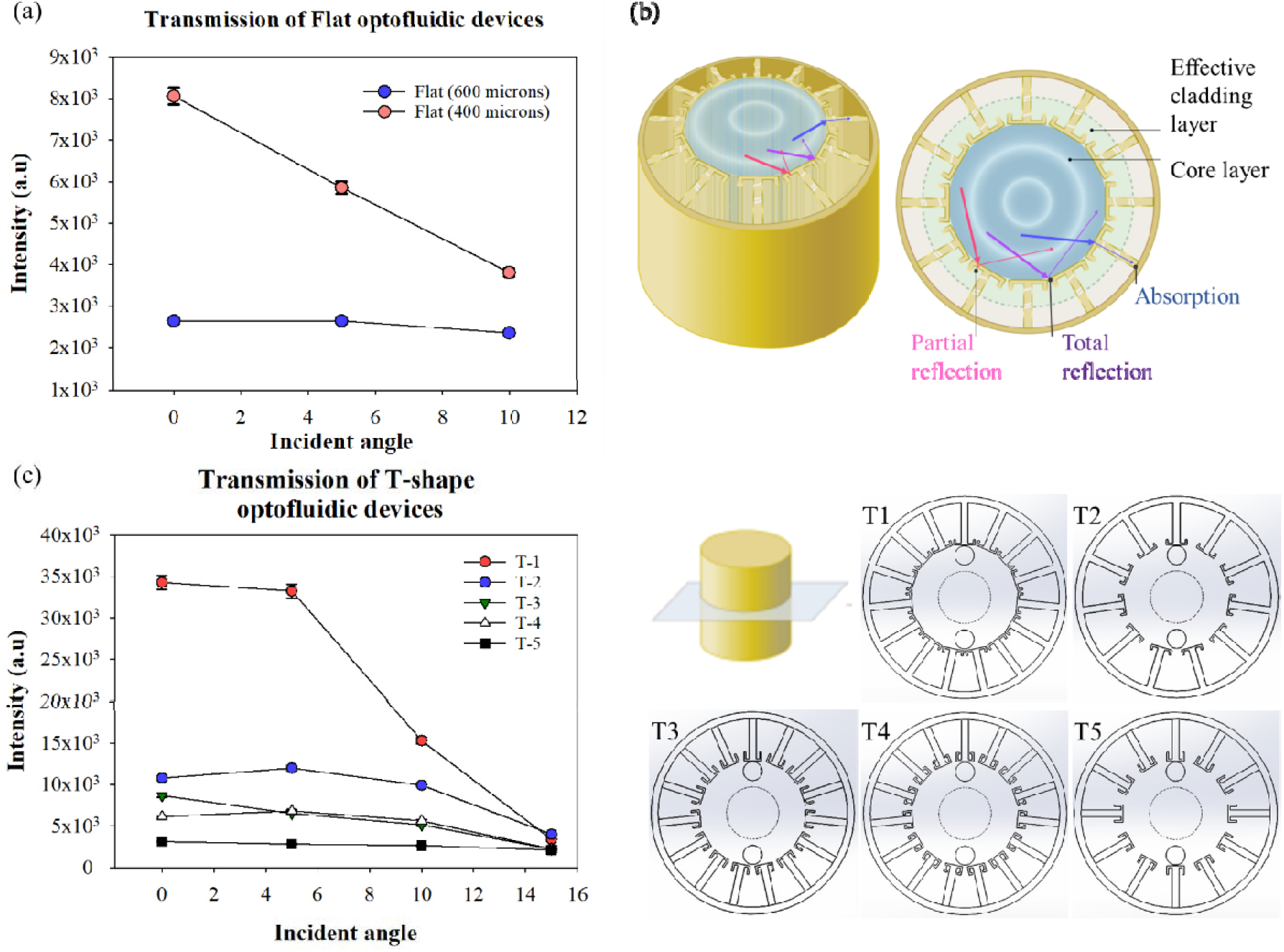
(a) Transmission measurements of a flat optofluidic chip with a solid cladding thickness of 400 μm and 600 μm, respectively. (b) The schematic of the solid/water/air interface to explain the loss mechanism. (c) Transmission measurements of various “T-shape” structures and incident angles as T-1 shows the best performance.

As shown in **Figure 3b**, the majority of the incident light beam (blue arrow) is absorbed by the solid wall while part of the light is reflected by the air cladding (purple arrow). Importantly, another part of the light is refracted and then reflected due to the combination of thin solid and air cladding (red arrow). Therefore, we demonstrated an ideal superhydrophobicity-based optofluidic configuration by minimizing the microstructure cap thickness and increasing the air cladding layer thickness, enabling an “air-mirror”^22^ at the solid/water/air interface. As a result, most of the light intensity will be reflected because of the thick air cladding layer.

The next factor investigated was how the cross-sectional shape of the liquid core can affect the light guiding as it can change the refraction of the light at the interface of solid/water/air. During the light propagation, the light rays will split into various refracted beams, and such a multipath phenomenon would lead to a nonlinear intensity. Ideally, a light guiding device consisting of a round core layer and a round cladding layer can better confine the refracted light rays in the center as well as more of the linear light intensity^20^. In **Figure 3c**, several “T-shape” devices were produced by varying the round shape smoothness at the interface between the core and cladding. Every device has nearly the same core layer volume as well as height of the microstructures. The solid fractions for T-1, T-3, and T-4 are similar at 37.5%, 38.4%, and 40%, respectively. While T-2 and T-4 have comparable solid fractions, which are 28.5% and 30.8% respectively.

T-1 has the best performance among the various structure designs. At a zero degree incident angle, the light transmission is at least 3.5 times better than the other designs. This is because the cross-section of T-1 is closest to a round shape and the width of the cap for T-1 is wider than the others except for T-2. The wider the T-cap is, the more “air-mirrors” created within the device. Therefore, more light can be reflected or partially reflected by the air cladding or air-thin solid mixed cladding layer. On the other hand, the core geometry of T-2’s cross-section is less round than T-1, meaning less light is reflected back to the liquid core regardless of the greater width. T-3, T-4, and T-5 also show more loss since they have narrower caps, resulting in less partial refraction. Ignoring the difference in the overhang lengths when compared to T-1 and T-2, the structures within T-3, T-4, and T-5 are longer than the other designs which may cause the greater intensity loss. Although the overhang structure provides the extra support necessary to hold the liquid, it still needs to be controlled at a reasonable thickness to avoid over increasing the solid substrate portion within the effective cladding layer. Lastly, T-5 has the lowest transmission due to a less circular cross-sectional geometry in addition to the longer length of the T-caps within the effective cladding thickness. These combined alterations to the microstructure design lead to a greater intensity loss caused by the solid substrate, and the narrower width of the T-caps.

**Figure 4** shows the application of the various designed optofluidic chips for fluorescence detection. Carboxyl quantum dots (size: 15-20 nm; Thermo Fisher Scientific) were prepared with a concentration ranging from 0.1 to 6.25 nM. The excitation wavelength was fixed at 405 nm with an incident angle of 18°. Three scans were averaged for each measurement with an integration time set at 100 ms. The uncorrected fluorescence spectra for the 1.6 nM quantum dots (211 μl) in different optofluidic chips are shown in **Figure 4a**, matching the trend from the transmission tests. The peak intensity for the “T-shape” device is ~2 times higher than the flat device, enabled by the TIR from air-cladding and air-thin solid mixed cladding. The integrated fluorescence intensity of various quantum dot concentrations are shown in **Figure 4b**, where the “T-shape” shows the best performance, and the measured intensity linearly increases with the concentration.

**Figure 4.**
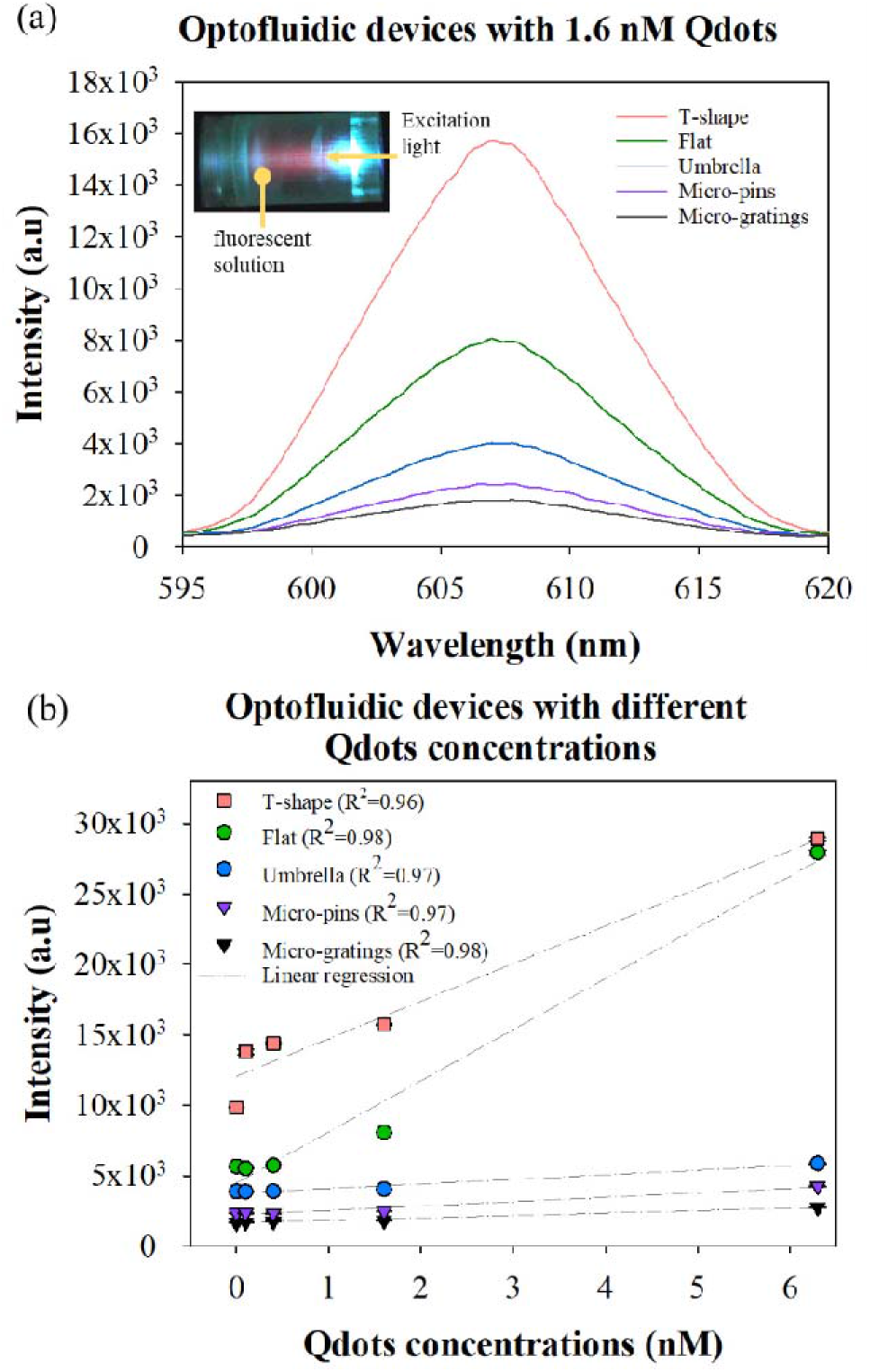
(a) Uncorrected fluorescence emission spectrum of various optofluidic platforms by inputting 1.6 nM Qdots. The inset is a photo of fluorescent solution (red) within the device. The light traveled from right to left. (b) Integrated fluorescence signal in the optofluidic platforms by increasing the quantum concentrations from 0 to 6.3 nM.

Lastly, we show that the optofluidic device can combine with a CRISPR-Cas12a system for detection of target DNA ^23,24^. As shown in **Figure 5a**, with the presence of target DNA, the single-stranded DNA probes are denatured by the CRISPR complex, thus leaving the quantum dots unconjugated in the solution^25^. On the other hand, without target DNA, the single-stranded DNA probes are intact, allowing the quantum dots to be captured by the magnetic beads and extracted from the necessary supernatant. The CRISPR-activated quantum dots remaining in the supernatant are then quantitatively evaluated by our optofluidics system. We set the quantum dot concentration at 1.68 nM and vary the DNA target concentration. As shown in **Figure 5b**, the integrated fluorescence intensity linearly increases with the increase in DNA target concentration from 0.1 to 1 nM. On the other hand, the sample without target input (NTC) has a lower signal than the 0.1 nM sample of target DNA strands.

**Figure 5.**
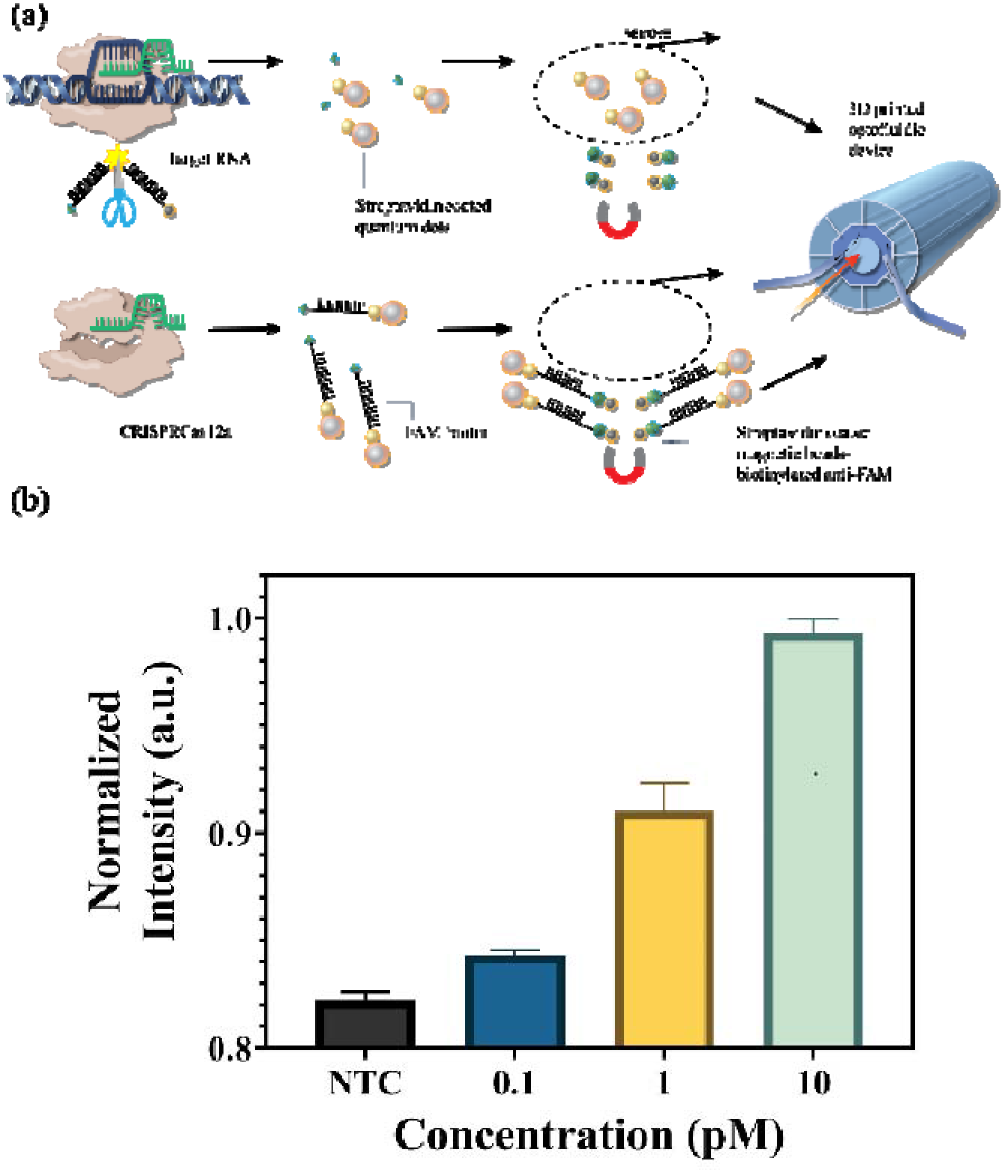
(a) The procedure of clustered regularly interspaced short palindromic repeat (CRISPR) in conjunction with-associated proteins (Cas) for simple and sensitive nucleic acid detection. (b) Fluorescence intensities of samples with DNA target input of 0.1, 1, and 10 nM. Negative control (without target) is labeled as NTC.

Different from solid core waveguides^26^, this fully-enclosed optofluidic system can efficiently collect the fluorescence signal of the CRISPR cleavage products in the liquid core, holding immense prospect for optogenetic therapy to deliver reagents locally or within specific tissues to treat various diseases such as sepsis^27^, cancer^28^, and neurological diseases^29^. Leveraging the high resolution of stereolithography, the “T-shape” structures can be fabricated with a smaller dimension for microsurgery while still providing efficient light transmission^30^. In the future, microvalves, micropumps, and LED light sources can also be 3-D printed and integrated with our optofluidics system as a single unit for sample loading, sensing, and treatment^31,32^.

## CONCLUSIONS

In summary, different designed geometries of superhydrophobic-based optofluidic devices were built by high resolution stereolithography techniques and examined via transmission and fluorescence measurements. The solid fraction within the range of theoretically required cladding thickness and the smoothness of the core’s cross-section were investigated to optimize our fully-enclosed optofluidic devices. Among all the designs, the T-shape optofluidic chip with a wide and thin T-cap, a short overhang length, and a round cross-section of the core shows the best performance with all the excitation wavelengths we tested, which agrees with our theoretical model. This T-shape optofluidic system was used to quantify a quantum dot-based CRISPR-Cas12 assay, thus establishing a novel technology capable of developing the next generation of CRISPR therapeutics for in vivo applications.

## AUTHOR INFORMATION

### Author Contributions

All authors have commented and given approval to the final version of the manuscript. ‡The author contributed to the design and implementation of the experiments. Yu Chang and Ke Du wrote the manuscript.

## ACKNOWLEDGMENT

The authors thank Sunghwan Bae (RIT) for the schematic design and Henry Yuqing for taking the SEM photos.

## ABBREVIATIONS

TIR: total internal reflection
PTFE: Polytetrafluoroethylene

